# Proteomic Analysis of Dorsal Root Ganglia in a Mouse Model of Paclitaxel-Induced Neuropathic Pain

**DOI:** 10.1101/2024.06.20.599888

**Authors:** Rania Hanna, Alexandru Graur, Patricia Sinclair, Bryan D. Mckiver, Paula D. Bos, M. Imad Damaj, Nadine Kabbani

**Affiliations:** Interdisciplinary Program in Neuroscience, George Mason University, Fairfax, VA 22030, USA; Department of Pharmacology & Toxicology, Virginia Commonwealth University, Richmond, VA 23298, USA; Department of Pathology, Massey Comprehensive Cancer Center, Virginia Commonwealth University School of Medicine, Richmond, VA, 23298

**Keywords:** neuropathy, chemotherapy, cancer, preclinical model, dorsal root ganglion

## Abstract

Paclitaxel is a chemotherapy drug widely used for the treatment of various cancers based on its ability to potently stabilize cellular microtubules and block division in cancer cells. Paclitaxel-based treatment, however, accumulates in peripheral system sensory neurons and leads to a high incidence rate (over 60%) of chemotherapy induced peripheral neuropathy. Using an established preclinical model of paclitaxel-induced peripheral neuropathy (PIPN), we examined proteomic changes in dorsal root ganglia (DRG) of adult male mice that were treated with paclitaxel (8 mg/kg, at 4 injections every other day) relative to vehicle-treated mice. High throughput proteomics based on liquid chromatography electrospray ionization mass spectrometry identified 165 significantly altered proteins in lumbar DRG. Gene ontology enrichment and bioinformatic analysis revealed an effect of paclitaxel on pathways for mitochondrial regulation, axonal function, and inflammatory purinergic signaling as well as microtubule activity. These findings provide insight into molecular mechanisms that can contribute to PIPN in patients.

## Introduction

Paclitaxel is a taxane class compound used as a cytotoxic chemotherapy agent in the treatment of various solid tumors including prostate, ovarian, breast cancer, bladder, esophageal, small and non-small-cell lung, pancreas, and melanoma [1,2]. Chemotherapy, with compounds such as paclitaxel, may interfere with a patient’s course of treatment due to the emergence of pain symptoms in over 50% of cases [3]. There is no effective mechanism for the prevention of chemotherapy associated neuropathy and conventional analgesics and other pain medications are generally ineffective in negative symptom management [4].

Paclitaxel acts by disrupting proliferation of cancer cells through binding with β-tubulin [5] and suppressing its depolymerization. However, paclitaxel can also target non-cancer cells leading to a number of side effects including skin cytotoxicity like rash and pruritus [6], and neuronal toxicity like peripheral neuropathy [7]. Studies show that paclitaxel treatment is also associated with changes in mitochondrial function, increased inflammation within dorsal root ganglia (DRG) and an increase in proapoptotic factors [7,8].

The DRG is a primary sensory module consisting of sensory neurons, satellite glia, and immune system cells such as macrophages and T cells [9]. Larger DRGs gives rise to myelinated axons that are involved in mechanoreception, whereas smaller DRGs give rise to unmyelinated axons, which are involved in mechanoreception,thermoreception, and nociception [10]. Paclitaxel has been shown to traverse the blood-nerve barrier surrounding the DRG and bind microtubules within neurons of DRG, causing damage to axons and nerve fibers [11]. Microtubule stabilization underlies axonal transport loss, which promotes axonal degeneration, leading to peripheral neuropathy [12]. Paclitaxel induced peripheral neuropathy (PIPN) is associated with damage to sensory axons and nerve fibers in humans and animal models [13,14].

Limited animal models exist to study molecular changes within sensory systems responding to paclitaxel exposure. Transcriptomics based studies of the DRG suggest an important role for paclitaxel in the activation of neuroinflammatory signaling within mice [14–16]. In this study we examined molecular mechanisms warrants studies at the level of the proteome. In this study, we examined molecular changes that may contribute to PIPN in a pre-clinical mouse model with heightened pain-like behaviors [17]. Specifically, the current proteomic analysis is conducted in mice that exhibit paclitaxel-induced mechanical hypersensitivity and cold hypersensitivity, as well as a decreased sensory nerve compound action potential (SNCAP) [17]. Our proteomic analysis aims to provide knowledge on mechanistic drivers of neuropathy during paclitaxel treatment.

## Methods

### Animals

The experiments were performed on 12-week-old male C57BL/6J mice (Strain #000664, The Jackson Laboratory, Bar Harbor, ME). Animals were housed in groups of 4 per cage with enriched environment, and maintained on a 12 hour light/dark cycle, at a 22°C room temperature with ad libitum access to food (global 18% protein chow diet; Envigo Teklad, Indianapolis, IN, USA) and water. DRG from 8 animals for both the experimental and control groups were used, with 3 mass spectrometry technical replicates completed.

Ethical statement: All animal experiments were performed under an approved IACUC protocol in the Division of Animal Research of Virginia Commonwealth University (Richmond, VA), accredited by the Association for Assessment and Accreditation of Laboratory Animal Care (AALAC). All efforts were made to reduce the number of animals used in this study and to ensure optimal conditions of well-being before, during and after each experiment. All mice were observed daily for general well-being and their weight was measured weekly. All behavioral experiments were performed during the light cycle and with the observer unaware of the experimental treatment of the animals.

### Drug treatment

Paclitaxel was purchased from VCU Health Pharmacy (Athenex, NDC 70860-200-50, Richmond, VA, USA) and dissolved in a 1:1:18 mixture of 200 proof ethanol, kolliphor, and distilled water (Sigma-Aldrich). Paclitaxel was administered at a dose of 8 mg/kg intraperitoneally (i.p.) every other day; 4 administrations completed one regimen. Control mice received a10 ml/kg i.p. injection of the diluent, with the same injection regimen. Seven days after drug treatment, mice were euthanized by decapitation. Lumbar (L4-L6) DRG tissues were collected, immediately frozen in liquid nitrogen, and stored at -80°C.

### Protein isolation

The DRG cytoplasmic and membrane fraction was obtained as described in [18]. Briefly, DRG specimens were combined with 500 μL of lysis buffer A (NaCl, HEPES, digitonin, hexylene glycol, protease, and phosphatase inhibitor cocktail) and disrupted for 5 seconds using a Dounce homogenizer. The resulting tissue suspension was processed with a QIAshredder homogenizer (Qiagen, 79656) and centrifuged at 500g for 10 minutes to filter the homogenate. The pellet was resuspended in 500 μL of lysis buffer A, incubated on a nutator for 10 minutes and centrifuged at 4000g for 10 minutes to isolate the cytosolic proteins.. The pellet was resuspended in 1 mL of lysis buffer B (NaCl, HEPES, Igepal, hexylene glycol, protease, and phosphatase inhibitor cocktail), incubated for 30-minutes on a nutator, then centrifuged at 6000g for 10 minutes to collect membrane bound proteins.. All centrifugation and incubation steps were performed at 4°C.

### Liquid-chromatography electrospray ionization mass spectrometry

Liquid-chromatography electrospray ionization mass spectrometry (LC-ESI MS/MS) was conducted in data-dependent acquisition (DDA) mode similar to previous studies [19,20]. Briefly, proteins were precipitated by incubaing for 5-minutes in acetone on ice followed by centrifugation. The protein pellet was denatured, reduced, and alkylated in 8 M urea, 1 M dithiothreitol, and 0.5 M iodoacetamide. Proteins were digested in trypsin (0.5 μg/μl) in 500 mM ammonium bicarbonate at 37°C for 5 hours and the fragments were desalted with C-18 ZipTips (Millipore), dehydrated in a SpeedVac for 18 mins, and reconstituted in 0.1% formic acid.

An Exploris Orbitrap 480 equipped with an EASY-nLC 1200 HPLC system (Thermo Fisher Scientific, Waltham, MA, USA) was used to conduct LC-ESI MS/MS analysis. Peptide separation was accomplished using a reverse-phase PepMap RSLC 75 μm i.d by 15 cm long with a 2 μm particle size C18 LC column (Thermo Fisher Scientific, Waltham, MA, USA). A solution of 80% acetonitrile and 0.1% formic acid was used for the peptide elution step at a flow rate of 300 nl/min. A full scan at 60,000 resolving power from 300 m/z to 1200 m/z was followed by peptide fragmentation with high-energy collision dissociation (HCD) at a normalized collision energy of 28%. EASY-IC filters were enabled for internal mass calibration, monoisotopic precursor selection, and dynamic exclusions (20 s). Data were recorded for peptide precursor ions with charge states ranging from +2 to +4. All samples were run in 3 technical replicates per condition.

### Proteomic and statistical analysis

Proteins were identified using the SEQUEST HT search engine within Proteome Discoverer v2.4 (Thermo Fisher Scientific, Waltham, MA, USA). Raw MS peptide spectra were compared to the NCBI mouse protein database using specific search engine parameters: mass tolerance for precursor ions of 2 ppm; mass tolerance for fragment ions of 0.05 Da; and a false discovery rate (FDR) cut-off value of 1% for reporting peptide spectrum matches (PSM) to the database.

Peptide abundance ratios were calculated by precursor ion quantification in Proteome Discoverer v2.4, using the vehicle control group as the denominator. Statistically significant abundance ratios with adjusted p-values < 0.05 were determined using the Student’s t-test. Analysis was performed on proteins with a quantifiable spectra signal profile observed in at least 2 of the 3 technical replicates. Proteomic data is deposited in the open access Figshare repository (https://doi.org/10.6084/m9.figshare.25905253).

### Bioinformatics

Gene ontology (GO) analysis was conducted in the Database for Annotation, Visualization, and Integrated Discovery (DAVID) [21,22]. The clustering of protein data was conducted using the Search Tool for the Retrieval of Interacting Genes/Proteins (STRING, v11.5) database using a Markov Cluster Algorithm (MCL) with an inflation parameter of 2 [23]. Data was processed, analyzed, and presented using Excel and the following tools: the R statistical software [24] and packages: ggplot2 [25], tidyverse [26]..

## Results

### DRG proteomics characterization in a mouse model of paclitaxel-induced neuropathy

Paclitaxel and other chemotherapy drugs elicit peripheral nerve fiber dysfunction and neurodegeneration that drives PIPN in humans [7]. Yet only a few rodent models exist for the study of PIPN including ours [28–30]. In recent studies, we have shown that paclitaxel administration drives altered behaviors including affective states and nociceptive responses such as mechanical allodynia and thermal hyperalgesia within mice [17,31–35]. To understand the mechanism that may underlie these sensory responses, we assessed proteomic changes from the DRG in response to a paclitaxel treatment. Eight adult male mice were treated with either 8 mg/kg body weight paclitaxel every other day for 4 days or 10 ml/kg of the vehicle diluent at the same injection regimen. Whole DRG were obtained from lumbar region L4-L6, an important site for sensory and pain processing [17]. Protein enrichment for cytosolic and membrane proteins was conducted prior to MS analysis. The overall workflow detailing the study is illustrated in **Fig. 1**.

**Figure 1.**
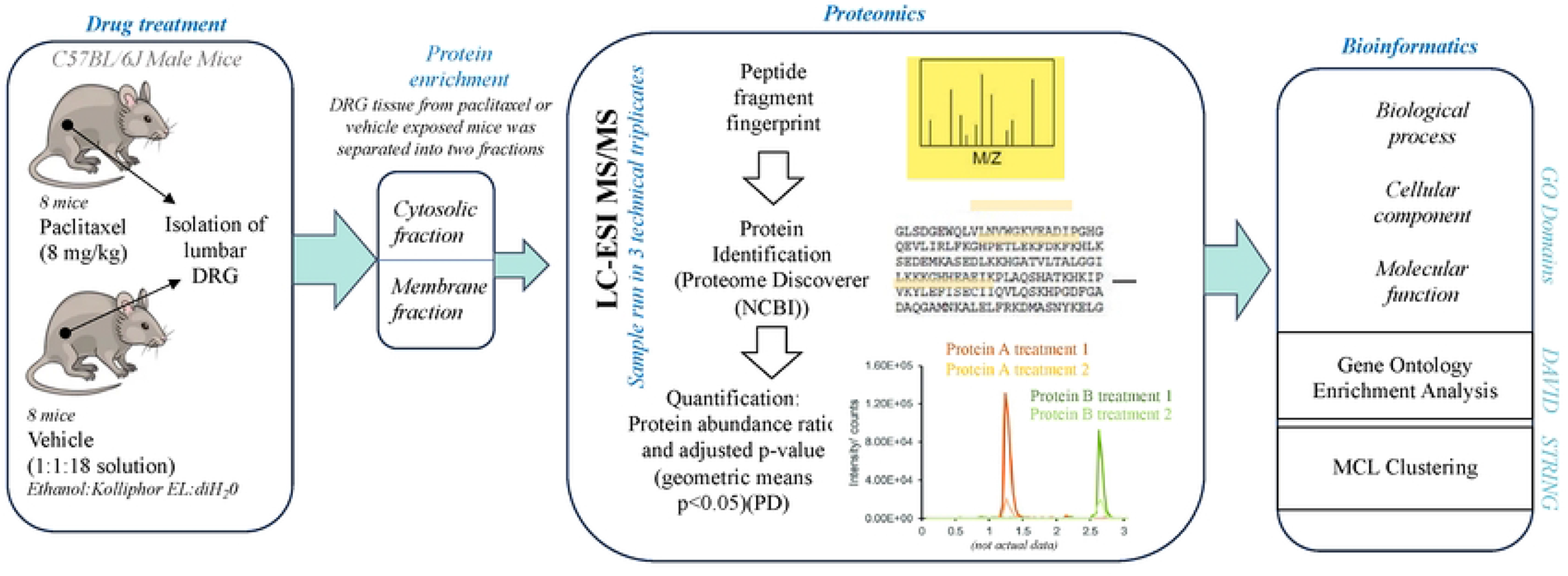
A workflow schematic showing the study design.

Proteomic analysis of DRG was conducted in order to compare changes in protein expression using label free peptide abundance analysis, as previously shown [19,20]. In these experiments fractions enriched for cytosolic and membrane obtained from the DRG of vehicle treated mice were used as the control. Using LC-ESI MS/MS, we identified a total of 2055 proteins within the cytosolic fraction, and 2676 within the membrane fraction (**Supplementary Table 1).** Statistical analysis of the abundance ratios indicates an effect of paclitaxel on 102 proteins within the membrane fraction and 63 proteins within the cytosol fraction. A complete list of significantly altered proteins from each fraction is presented in **Supplementary Tables 2-5**. Of the significantly altered proteins, 36 proteins are upregulated while 27 proteins are downregulated in the cytosolic fraction (**Fig. 2A**) (**Supplementary Table 4, 5**). In addition, 28 proteins are upregulated while 74 are downregulated in the membrane fraction (**Fig. 2B**) (**Supplementary Table 2, 3**).

**Figure 2.**
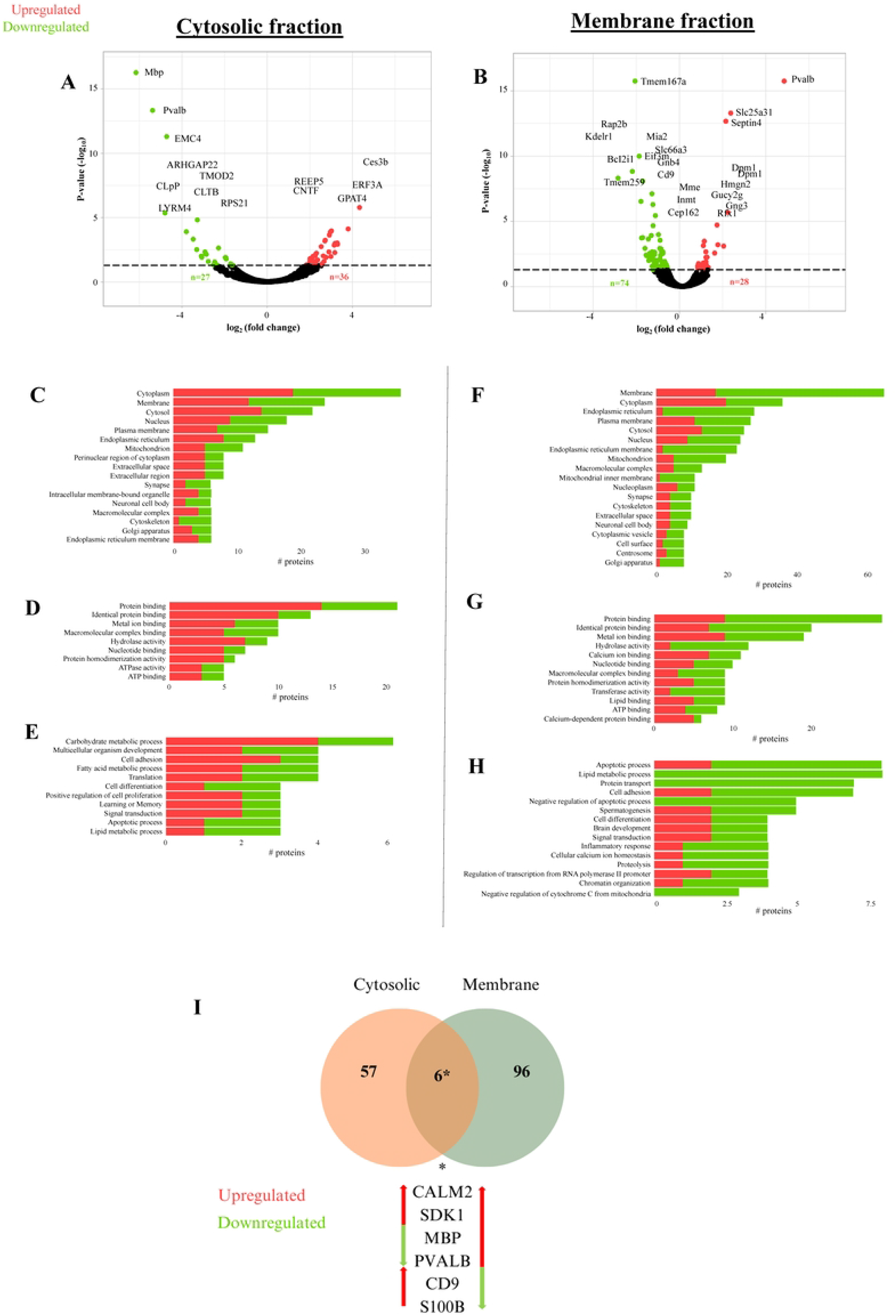
Effect of paclitaxel on the DRG proteome. Volcano plot distribution of significantly altered proteins within the DRG. A) Cytosolic fraction. B) Membrane fraction. GO terms associated with the significantly altered proteins DRG of paclitaxel treated mice relative to controls. Cytosolic fraction cellular component (C), molecular function (D), biological process. (E). Membrane fraction cellular component (F), molecular function (G), biological process (H).

We compared our proteomic dataset to published proteomic studies from mouse DRG [36–38]. 88% of proteins identified within our DRG analysis also appear in the DRG proteome of adult mice within pain related studies [38]. These overlapping DRG proteins are shown in **Supplementary** Fig. 1 and **Supplementary Table 6** [39]. A GO analysis of the significantly altered proteins indicates enrichment of cellular components, molecular functions and biological processes across cytosolic and membrane fraction proteins in the DRG of paclitaxel-treated mice (**Fig. 2**). Specifically, cytosolic fraction results indicate an effect of paclitaxel on carbohydrate metabolic processes, lipid metabolism, ATPase function, cytoskeletal intermediate filament and cell adhesion, and cell proliferation as well as apoptosis (**Fig. 2C-E**). Membrane fraction results show an effect of paclitaxel on ATP activity, cell adhesion, cytoskeletal organization, protein transport, and calcium and lipid signaling (**Fig. 2F-H**). GO results appeared to slightly differ however between the two fraction protein results with cytosolic proteomes highlighting an increase in carbohydrate metabolic processes, while the membrane fraction underscores a strong effect of paclitaxel on attenuated lipid metabolism. GO informatic analysis of the DRG proteome shows an effect of paclitaxel on several important metabolic pathways as well as signaling and structural functions within the DRG.

### An effect of paclitaxel on DRG energy, cytoskeletal regulation, and sensory axon signaling

We used DAVID [22,27] enrichment analysis of GO terms that are associated with significantly altered proteins in both cytosolic and membrane fractions to compare the effect of paclitaxel on DRG proteome adaptation. As shown in **Tables 1 and 2**, enrichment analysis confirms representation of membrane and cytosolic terms within the membrane and cytosolic fractions, respectively. Additionally, enrichment analysis of the membrane fraction indicates a high representation of membrane proteins from various intracellular organelles including the mitochondria as well as the endoplasmic reticulum (ER). Membrane associated proteins are also involved in lipid, metabolic and calcium cell signaling (**Table 2**). We assessed for overlap in significantly altered proteins across the membrane and cytosolic fractions. A subset of significantly altered proteins appeared in both membrane and cytosolic fractions results (**Fig. 2I**). These proteins include cluster of differentiation 9 (CD9), myelin basic protein (MBP), calcium binding proteins S100B, parvalbumin (Pvalb) and calmodulin, as well as the cell adhesion molecule SDK1. An analysis of the effect of paclitaxel reveals that some of these overlapping proteins are differentially regulated by the drug treatment condition (**Fig. 2I**). These findings suggest an effect of paclitaxel on specific proteins within the DRG.

**Table 1:** DAVID enrichment terms and their associated proteins within the cytosolic fraction.

| **GO category** | **Term** | **Count** | **Proteins** | **Adjusted p-value** |
| --- | --- | --- | --- | --- |
| Cellular component | Cytoplasm | 36 | YWHAЕ, MTPN, CNTF, TAGLN, GSPT1, CST3, ARHG AP22, HEBP2, MBP, PTGDS, GET3, EIF4E, PHM1, APC2, TMP3, TMOD2, CDC42BPB, TBCA, S100B, TSN, KIF27, PPP1CA, CARHSP1, RPS28, PSMA4, STK24, NACA, SLMAP, GPD1, G6PDX, GNB3, NAA15, GPD1L, CALM2, RPS21, PVALB | 0.009819583 |
| Molecular function | Macromolecular complex binding | 10 | YWHAЕ, CNTF, TTR, VAPA, CDC42BPB, ATP5F1D, TSN, ATP5F1E, PVALB, PPP1CA | 0.027691179 |

**Table 2:** DAVID enrichment terms and their associated proteins within the membrane fraction.

| GO category | Term | Count | Proteins | Adjusted P-value |
| --- | --- | --- | --- | --- |
| Cellular component | Membrane | 64 | LGALS3BP, CLIC4, GOLT1B, KDELR1, ICAM2, MIA2, ATP2A1, FNBP1L, COX7C, TMEM186, CLGN, NDST3, ANXA6, SLC36A1, ENTPD1, MME, SIGMAR1, TAP1, DYNLL1, TLCD3B, MFSD10, DPM1, TMEM256, RAP2B, STIM1, DAD1, GORASP2, TMEM259, NDUFS4, S100A6, CD47, S100A9, CERS2, CDS1, RTN3, STOML2, SLC66A3, MGST3, PON2, DERL1, SLC1A4, ASAP2, HIGD1A, GUCY2G, ZMPSTE24, MTDH, PTDSS2, GNG3, MBP, CD99, MPDUL1, S100A10, NDUFA3, SURF4, ERLIN1, SDK1, TSPAN18, CD9, MAN2B1, ESYT2, SLC25A31, CALM2, TOMM5, BCL2L1, CDS2 | 0.00004 |
| Cellular component | Endoplasmic reticulum | 28 | CDS1, RTN3, GOLT1B, KDELR1, MGST3, DERL1, MIA2, ATPSA1, CLGN, MTDH, ZMPSTE24, PTDSS2, SLC36A1, RPS7, SURF4, SIGMAR1, ERLIN1, TAP1, TLCD3B, DPM1, STIM1, DAD1, GORASP2, TMEM259, ESYT2, BCL2L1, CERS2, CDS2 | 0.00000035 |
| Cellular component | Endoplasmic reticulum membrane | 23 | CDS1, RTN3, KDELR1, SURF4, SIGMAR1, ERLIN1, DERL1, TAP1, MIA2, ATP2A1, TLCD3B, CLGN, MTDH, ZMPSTE24, DPM1, PTDSS2, STIM1, DAD1, GORASP2, TMEM259, ESYT2, CERS2, CDS2 | .000000294 |
| Cellular component | Mitochondrion | 20 | STOML2, CLIC4, NDUFA3, MGST3, TAP1, ATP2A1, SEPTIN4, DYNLL1, HIGD1A, COX7C, TMEM186, TXN2, NDUFS4, ANXA6, ACOT2, PPIF, | 0.030463258 |
|  |  |  | PMPCB, SLC25A31, TOMM5, BCL2L1 |  |
| Cellular component | Macromolecular complex | 13 | PRPS1, RPS7, GOLT1B, ERLINI, HIGD1A, ZMPSTE24, STIM1, ANXA6, GNB4, CD9, MBP, CALM2, PVALB | 0.045210595 |
| Cellular component | Mitochondrial inner membrane | 11 | STOML2, TMEM256, NDUFA3, NDUFS4, PPIF, PMPCB, SLC25A31, HIGD1A, COX7C, TMEM186, BCL2L1 | 0.0093664559 |
| Molecular function | Lipid binding | 9 | STOML2, FABP4, APOH, ERLINI, ANXA6, PMP2, ESYT2, MBP, FNBP1L | 0.031904593 |
| Molecular function | Calcium-dependent protein binding | 6 | ANXA6, S100A6, S100B, CALM2, S100A9, S100A10 | 0.016979225 |

Proteomic analysis can also offer insight into drug-treatment associated adaptations within protein-protein interaction (PPI) networks within target tissue and cell types [40].We used a Markov cluster (MCL) analysis to define PPI networks within the DRG proteome consisting of all significantly upregulated and downregulated proteins from membrane and cytosolic fractions. MCL analysis is represented by an integrated PPI network based on the identity of the significantly altered proteins in response to paclitaxel treatment (**Fig. 3**). Within the PPI network we identified 11 high confidence protein clusters. The largest PPI cluster (Cluster 1) was found to contain 14 proteins with 47 connections yielding a significant PPI connection (p < 1.0 × 10^−16^). It consists of mitochondrial proteins and is positioned centrally within the integrated PPI map (**Fig. 3**). Cluster 1 had an average local clustering coefficient (ALCC) of 0.755 with analysis revealing downregulation in this mitochondrial protein network (**Fig. 4**). The second cluster (Cluster 2) consists of 12 ribosomal proteins that also appear predominantly downregulated within the DRG in response to paclitaxel (**Fig. 4**). Additional clusters identified within PPI analysis include cell structural and motility regulators that include cytoskeletal elements (e.g., Clusters 8, 10, 11) and protein clusters involved in biochemical pathways including nitrogen and pyridine metabolism (Clusters 6 and 7).

**Figure 3.**
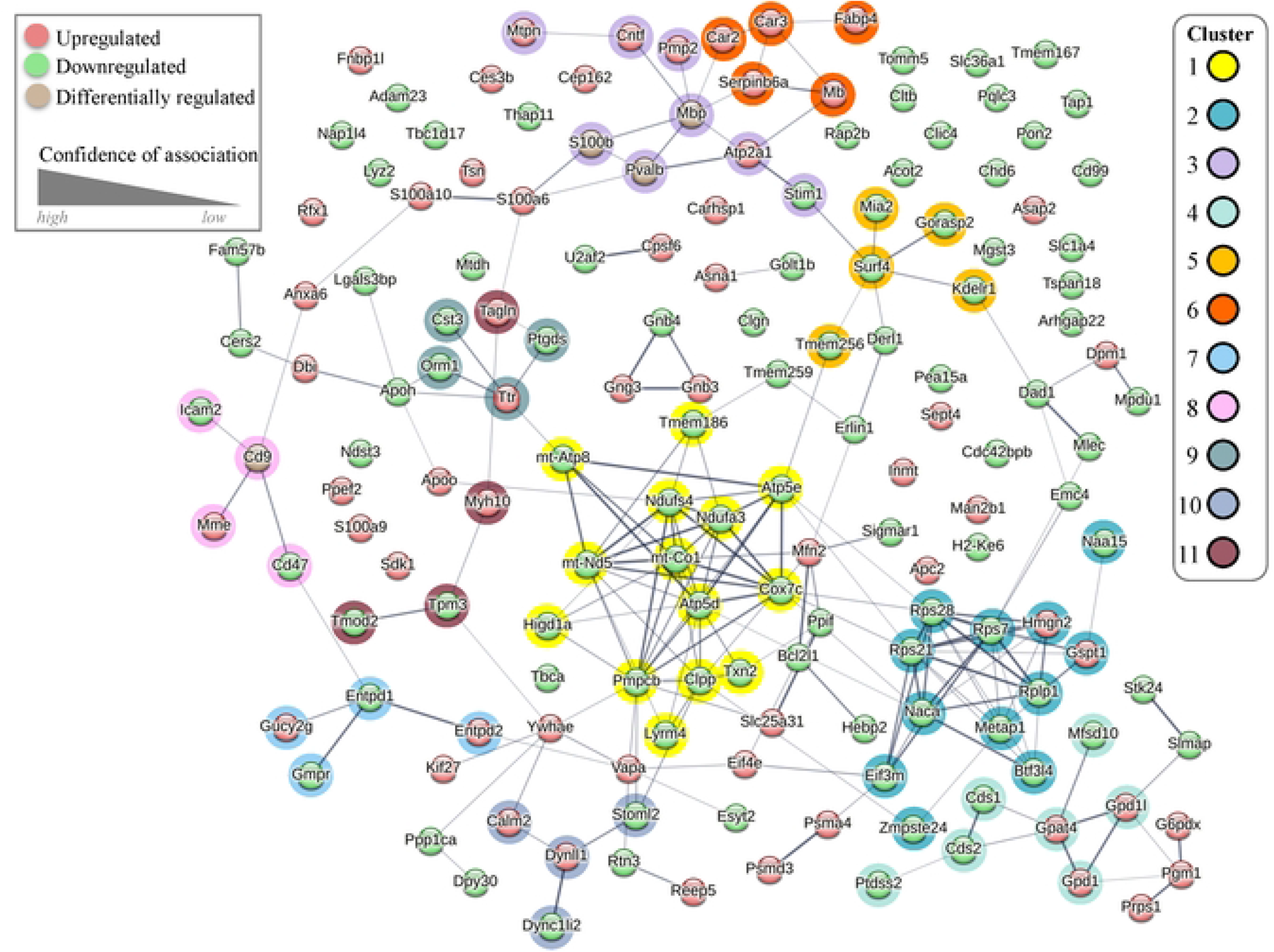
A protein-protein interaction (PPI) map of the paclitaxel associated DRG proteome. STRING networks showing PPI amongst significantly altered proteins. Protein clusters within the network are highlighted by color. The thickness of the connection indicates the degree of confidence between node associations while color indicates whether the protein is increased (red), decreased (green) or differentially altered (light brown) in the paclitaxel condition. The STRING network is based on an MCL algorithm with an inflation parameter of 2 used to identify up to 11 cluster groups.

**Figure 4.**
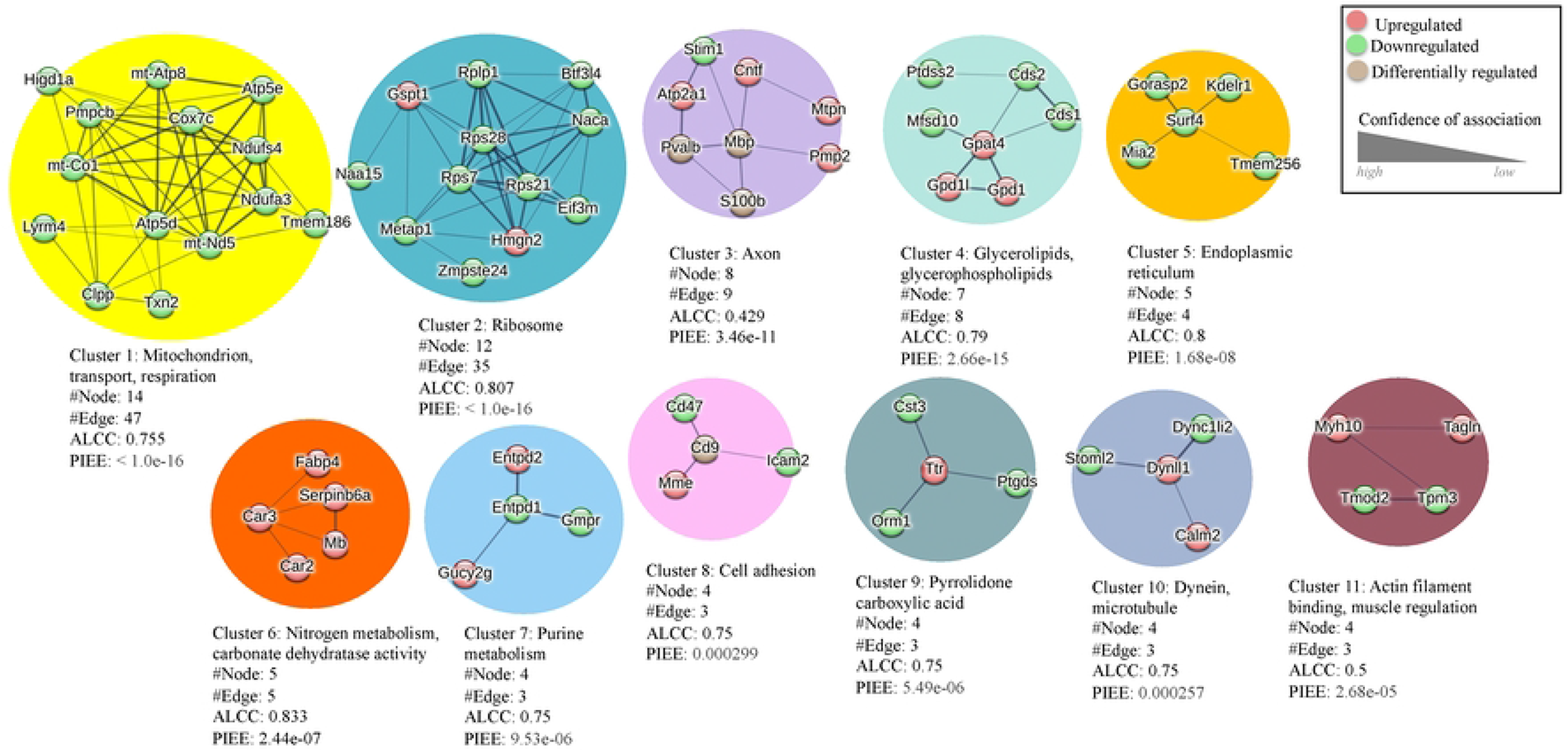
Primary cluster groups within the PPI network. An enrichment tag analysis showing the 11 cluster groups within STRING analysis of the paclitaxel associated DRG proteome.

## Discussion

We used a proteomics research strategy to examine molecular changes within the DRG in an established mouse model of paclitaxel induced neuropathy [17,41]. In recent studies, multi-OMIC methods such as metabolomic, proteomic, and phospho-proteomic analyses were used to identify molecular changes in the DRG during diabetic neuropathy in humans [42]. Our study expands this methodology and assesses the specific impact of paclitaxel on the DRG proteome within a PIPN mouse model. Overall, our proteomic results highlight putative disruption to mitochondrial components and related metabolic processes within the DRG during paclitaxel administration, similar to studies that indicate that paclitaxel causes mitochondrial damage, leading to other issues, including axonal damage [43–45]. In addition, our findings support the role of paclitaxel in the remodeling of intracellular compartments such as the ER, Golgi, and the cytoskeleton through changes in actin filament and dynein microtubule proteins [44–46]. These results provide evidence on the role of paclitaxel in nervous tissue, and shed light on the potential impact of chemotherapy on DRG associated sensory and neuroimmune activity [47,48].

Paclitaxel binds to the microtubule cytoskeleton in both cancer and non-cancer cells impairing cell division as well as other general functions. Studies in rodent models show that paclitaxel treatment can also lead to the demyelination and degeneration of axons in nervous tissue [49]. Paclitaxel treatment is also shown to disrupt microtubule transport along long axons thereby impairing protein and organelle trafficking within the nervous system [50,51]. In particular, paclitaxel’s actions appear to correlate with deficits in mitochondrial trafficking in axons both *in vitro* as well as *in vivo* [50,51] A schematic model of our proteomic findings on DRG function is presented in **Fig. 5**. In this model, sensory deficits are suggested to arise as a function of protein changes within multiple cell types and processes in the DRG. In addition, paclitaxel associated proteomic changes may impact sensory signaling outside of the DRG through interactions of sensory axons with target tissue (e.g. epidermis) thereby contributing to long-term pain sensitivity.

**Figure 5.**
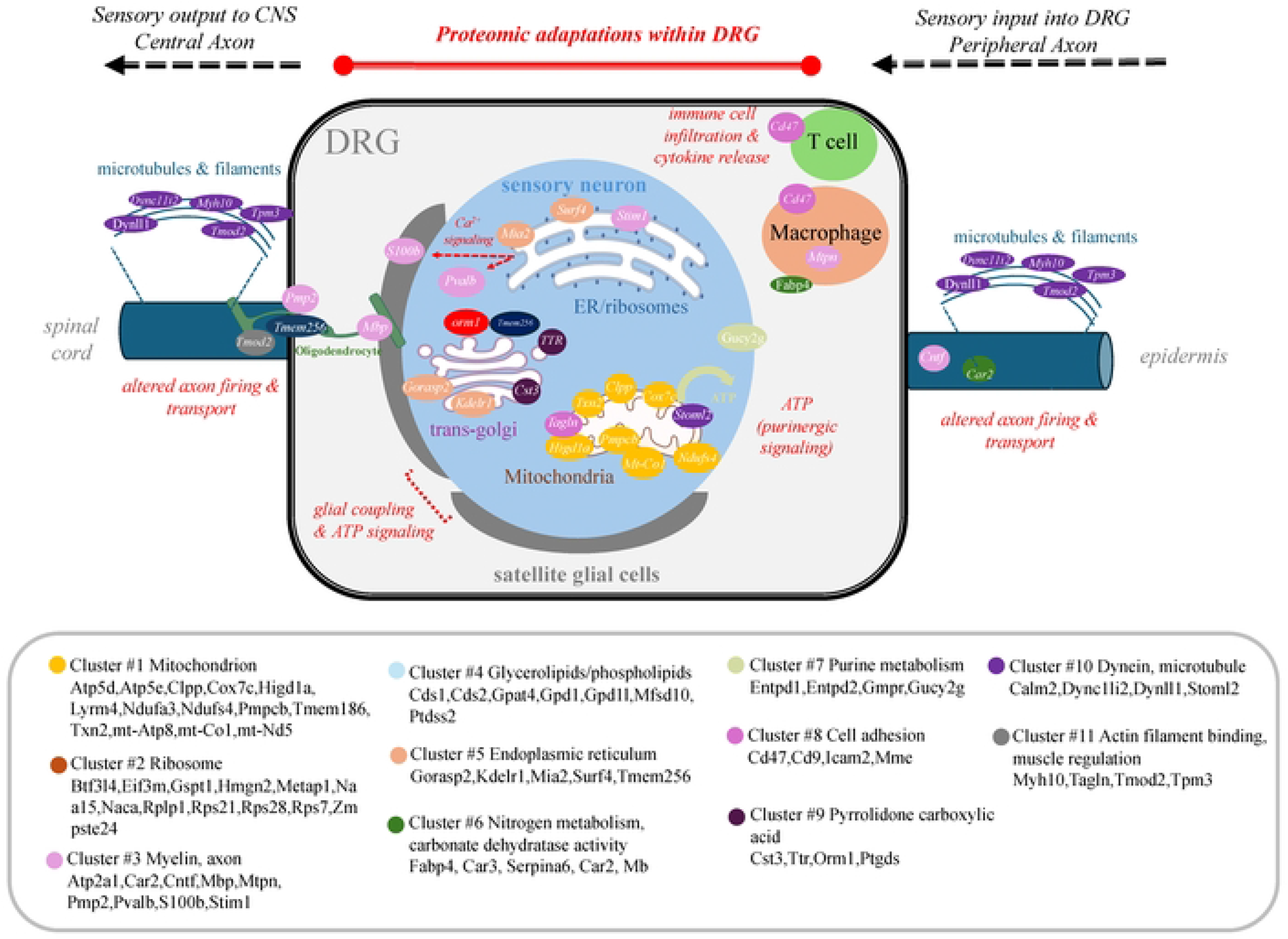
A pathway and signaling model suggestive of the involvement of altered proteins within the DRG.

Our model is consistent with evidence on the involvement of DRG connectivity in chemotherapy associated neuropathy [52]. The DRG contains various sensory neurons, which serve in the processing of sensory information between periphery and CNS. These sensory neurons are encapsulated by a satellite glial cell (SGC) network that modulates neuronal responses to nociception [53]. In addition, immune cells can enter the DRG under heightened inflammatory conditions [54]. Our findings support the involvement of both neurons and glia in the effect of paclitaxel within the DRG [17] as evidenced by changes in the expression of cell type proteins. For example, parvalbumin (Pvalb), a calcium binding protein that identifies a subpopulation of proprioceptive DRG neurons and has been shown to be associated with peripheral nerve injury [55], is found to be significantly decreased by paclitaxel. S100b on the other hand, a calcium binding protein expressed in glial cells of the DRG [56], is also found to be decreased within the paclitaxel treatment groups. These findings point to an effect of paclitaxel on various cell types within the DRG and suggest that cell specific proteome analysis is an important future direction in mechanistic study of changes during PIPN.

Bioinformatic analysis of paclitaxel associated proteomic changes within the DRG defines 11 functional network clusters with components that are already known to contribute to pain responses in animal models [57–59]. Our results show proteins across various clusters that contribute to enhanced inflammatory responses within the DRG. For example, changes in the expression of immune cell markers including orosomucoid 1 protein (Orm1), an acute phase inflammatory response protein, and CD47 support evidence on increased neuroimmune activity within the DRG during paclitaxel treatment [60]. Changes in immune cell activity may also contribute to a loss in oligodendrocyte function including myelin production during neuropathic pain [61]. In support of this, our results show that myelin basic protein (Mbp) is significantly altered within the DRG in response to paclitaxel.

Non-neuronal cells and particularly SGCs are increasingly recognized for their role in the etiology of neuropathic pain. SGC are coupled via gap junctions that are modified by nerve injury as well as neuroinflammation [54]. Chemotherapeutic drugs, such as oxaliplatin or paclitaxel, have been shown to alter the number of gap junctions between SGCs leading to increased electrical coupling and calcium oscillatory activity within the DRG [62,63]. In addition, SGC transmit signals bidirectionally between neuron and glia through purinergic (ATP) signaling [64]. Our data shows that Cluster 7 is enriched in purine metabolism proteins that are important for ATP signaling. Changes in ATP sensitivity within the DRG have been previously demonstrated in a mouse pain model with mechanical hypersensitivity and DRG sensory neuron hyperexcitability [45,65,66]. An analysis of differentially expressed proteins points to an enrichment of ATP regulatory proteins that are also associated with mitochondrial activity. The largest of our clusters (Cluster 1) represents mitochondrial proteins including cytochrome C oxidase subunit 7c (Cox7c), mitochondrial encoded cytochrome C oxidase 1 (mt-Co1), thioredoxin 2 (Txn2), and hypoxia-inducible domain family member 1a (Higd1a). Mitochondrial dysfunction is a common feature of neuropathic pain including PIPN [42]. Changes in mitochondrial proteins can impact multiple cell types within the DRG and may influence processes such as Schwann cell myelination [67].

Our findings involve nitrogen metabolism (Cluster 6), which fits in with existing research of the role nitrogen-related compounds have in neuropathic pain. Namely, myoglobin (Mb) has been shown to be involved in pain processes [68]. Within this pathway, nitric oxide (NO) is important in a number of signaling pathways, including immune regulation, neuronal survival, and synaptic plasticity [69], with an overproduction of NO associated with neuropathic pain [70]. Molecules associated with nitrogen metabolism have been shown to modulate sensory neuron excitability and nociceptive effects [71]. Within Cluster 6, we find that the fatty acid binding protein 4 (FABP4) is significantly increased within the DRG in response to paclitaxel administration.

FABP4 is expressed in adipocytes as well as immune cells such as macrophages and participates in lipid signaling as well as nitrogen metabolism. Studies show that FABP4 can contribute to neuroinflammation [72].

Our findings provide insight on mechanisms that can contribute to pain responses following paclitaxel treatment within our PIPN mouse model. Our proteomic results are consistent with earlier studies in this mouse model that show an effect of paclitaxel on mechanical and thermal sensitivity . This work is also similar to proteomic studies of paclitaxel associated neuropathy within the rat DRG [41]. Understanding the mechanisms that contribute to PIPN is an important research direction that can be leveraged through combining several OMIC technologies including RNA sequencing (RNAseq) and proteomics withing these established preclinical models.

## Acknowledgements

This work was funded by R01CA219637 to MID.

## Data availability

Proteomic data generated during this study is deposited in the online open access Figshare repository (https://doi.org/10.6084/m9.figshare.25905253).

## Notes

### Competing Interest Statement

The authors have declared no competing interest.

## References

1. Belani CP. Paclitaxel and docetaxel combinations in non-small cell lung cancer. Chest. 2000;117: 144S–151S. doi:10.1378/chest.117.4_suppl_1.144s

2. Nathan FE, Berd D, Sato T, Mastrangelo MJ. Paclitaxel and tamoxifen: An active regimen for patients with metastatic melanoma. Cancer. 2000;88: 79–87. doi:10.1002/(sici)1097-0142(20000101)88:1<79::aid-cncr12>3.0.co;2-l

3. Polomano RC, Farrar JT. Pain and neuropathy in cancer survivors. Surgery, radiation, and chemotherapy can cause pain; research could improve its detection and treatment. Am J Nurs. 2006;106: 39–47. doi:10.1097/00000446-200603003-00015

4. Kawashiri T, Inoue M, Mori K, Kobayashi D, Mine K, Ushio S, et al. Preclinical and Clinical Evidence of Therapeutic Agents for Paclitaxel-Induced Peripheral Neuropathy. Int J Mol Sci. 2021;22: 8733. doi:10.3390/ijms22168733

5. Risinger AL, Riffle SM, Lopus M, Jordan MA, Wilson L, Mooberry SL. The taccalonolides and paclitaxel cause distinct effects on microtubule dynamics and aster formation. Mol Cancer. 2014;13: 41. doi:10.1186/1476-4598-13-41

6. Li H-B, Wang W-H, Wang Z-Y. A meta-analysis of the incidence and risk of skin toxicity with nab-paclitaxel and paclitaxel in cancer treatment. Am J Transl Res. 2023;15: 4279– 4290.

7. Klein I, Lehmann HC. Pathomechanisms of Paclitaxel-Induced Peripheral Neuropathy. Toxics. 2021;9: 229. doi:10.3390/toxics9100229

8. Bhalla KN. Microtubule-targeted anticancer agents and apoptosis. Oncogene. 2003;22: 9075–9086. doi:10.1038/sj.onc.1207233

9. 9. Ahimsadasan N, Reddy V, Khan Suheb MZ, Kumar A. Neuroanatomy, Dorsal Root Ganglion. StatPearls. Treasure Island (FL): StatPearls Publishing; 2024. Available: http://www.ncbi.nlm.nih.gov/books/NBK532291/

10. Cervero F, Laird JMA. Mechanisms of touch-evoked pain (allodynia): a new model. Pain. 1996;68: 13–23. doi:10.1016/S0304-3959(96)03165-X

11. Gornstein EL, Schwarz TL. Neurotoxic mechanisms of paclitaxel are local to the distal axon and independent of transport defects. Exp Neurol. 2017;288: 153–166. doi:10.1016/j.expneurol.2016.11.015

12. Fukuda Y, Li Y, Segal RA. A Mechanistic Understanding of Axon Degeneration in Chemotherapy-Induced Peripheral Neuropathy. Front Neurosci. 2017;11: 481. doi:10.3389/fnins.2017.00481

13. Vermeer CJC, Hiensch AE, Cleenewerk L, May AM, Eijkelkamp N. Neuro-immune interactions in paclitaxel-induced peripheral neuropathy. Acta Oncol. 2021;60: 1369–1382. doi:10.1080/0284186X.2021.1954241

14. Sun W, Yang S, Wu S, Ba X, Xiong D, Xiao L, et al. Transcriptome analysis reveals dysregulation of inflammatory and neuronal function in dorsal root ganglion of paclitaxel-induced peripheral neuropathy rats. Mol Pain. 2022;19: 17448069221106167. doi:10.1177/17448069221106167

15. Kim HK, Lee S-Y, Koike N, Kim E, Wirianto M, Burish MJ, et al. Circadian regulation of chemotherapy-induced peripheral neuropathic pain and the underlying transcriptomic landscape. Sci Rep. 2020;10: 13844. doi:10.1038/s41598-020-70757-w

16. Housley SN, Nardelli P, Carrasco DI, Rotterman TM, Pfahl E, Matyunina LV, et al. Cancer Exacerbates Chemotherapy-Induced Sensory Neuropathy. Cancer Res. 2020;80: 2940– 2955. doi:10.1158/0008-5472.CAN-19-2331

17. Caillaud M, Patel NH, White A, Wood M, Contreras KM, Toma W, et al. Targeting Peroxisome Proliferator-Activated Receptor-α (PPAR-α) to Reduce Paclitaxel-Induced Peripheral Neuropathy. Brain Behav Immun. 2021;93: 172–185. doi:10.1016/j.bbi.2021.01.004

18. Baghirova S, Hughes BG, Hendzel MJ, Schulz R. Sequential fractionation and isolation of subcellular proteins from tissue or cultured cells. MethodsX. 2015;2: 440–445. doi:10.1016/j.mex.2015.11.001

19. Graur A, Sinclair P, Schneeweis AK, Pak DT, Kabbani N. The human acetylcholinesterase C-terminal T30 peptide activates neuronal growth through alpha 7 nicotinic acetylcholine receptors and the mTOR pathway. Sci Rep. 2023;13: 11434. doi:10.1038/s41598-023-38637-1

20. Sinclair P, Kabbani N. Nicotinic receptor components of amyloid beta 42 proteome regulation in human neural cells. Araki W, editor. PLoS ONE. 2022;17: e0270479. doi:10.1371/journal.pone.0270479

21. Sherman BT, Hao M, Qiu J, Jiao X, Baseler MW, Lane HC, et al. DAVID: a web server for functional enrichment analysis and functional annotation of gene lists (2021 update). Nucleic Acids Res. 2022;50: W216–W221. doi:10.1093/nar/gkac194

22. Huang DW, Sherman BT, Lempicki RA. Systematic and integrative analysis of large gene lists using DAVID bioinformatics resources. Nat Protoc. 2009;4: 44–57. doi:10.1038/nprot.2008.211

23. Szklarczyk D, Kirsch R, Koutrouli M, Nastou K, Mehryary F, Hachilif R, et al. The STRING database in 2023: protein–protein association networks and functional enrichment analyses for any sequenced genome of interest. Nucleic Acids Research. 2023;51: D638– D646. doi:10.1093/nar/gkac1000

24. R Core Team. R: A language and environment for statistical computing. Vienna, Austria: R Foundation for Statistical Computing; 2021. Available: https://www.R-project.org/

25. 25. Wickham H. ggplot2: Elegant graphics for data analysis. Springer-Verlag, New York; 2016. Available: https://ggplot2.tidyverse.org

26. Wickham H, Averick M, Bryan J, Chang W, McGowan L, François R, et al. Welcome to the Tidyverse. JOSS. 2019;4: 1686. doi:10.21105/joss.01686

27. Sherman BT, Hao M, Qiu J, Jiao X, Baseler MW, Lane HC, et al. DAVID: a web server for functional enrichment analysis and functional annotation of gene lists (2021 update). Nucleic Acids Research. 2022;50: W216–W221. doi:10.1093/nar/gkac194

28. Höke A, Ray M. Rodent Models of Chemotherapy-Induced Peripheral Neuropathy. ILAR Journal. 2014;54: 273–281. doi:10.1093/ilar/ilt053

29. Toma W, Caillaud M, Patel NH, Tran TH, Donvito G, Roberts J, et al. N-acylethanolamine-hydrolysing acid amidase: A new potential target to treat paclitaxel-induced neuropathy. European Journal of Pain. 2021;25: 1367–1380. doi:10.1002/ejp.1758

30. Toma W, Kyte SL, Bagdas D, Alkhlaif Y, Alsharari SD, Lichtman AH, et al. Effects of paclitaxel on the development of neuropathy and affective behaviors in the mouse. Neuropharmacology. 2017;117: 305–315. doi:10.1016/j.neuropharm.2017.02.020

31. Curry ZA, Wilkerson JL, Bagdas D, Kyte SL, Patel N, Donvito G, et al. Monoacylglycerol Lipase Inhibitors Reverse Paclitaxel-Induced Nociceptive Behavior and Proinflammatory Markers in a Mouse Model of Chemotherapy-Induced Neuropathy. J Pharmacol Exp Ther. 2018;366: 169–183. doi:10.1124/jpet.117.245704

32. Contreras KM, Caillaud M, Neddenriep B, Bagdas D, Roberts JL, Ulker E, et al. Deficit in voluntary wheel running in chronic inflammatory and neuropathic pain models in mice: Impact of sex and genotype. Behavioural Brain Research. 2021;399: 113009. doi:10.1016/j.bbr.2020.113009

33. Kyte SL, Toma W, Bagdas D, Meade JA, Schurman LD, Lichtman AH, et al. Nicotine Prevents and Reverses Paclitaxel-Induced Mechanical Allodynia in a Mouse Model of CIPN. J Pharmacol Exp Ther. 2018;364: 110–119. doi:10.1124/jpet.117.243972

34. Meade JA, Alkhlaif Y, Contreras KM, Obeng S, Toma W, Sim-Selley LJ, et al. Kappa opioid receptors mediate an initial aversive component of paclitaxel-induced neuropathy. Psychopharmacology (Berl). 2020;237: 2777–2793. doi:10.1007/s00213-020-05572-2

35. Xu L, Chen X, Wang L, Han J, Wang Q, Liu S, et al. Paclitaxel combined with platinum (PTX) versus fluorouracil combined with cisplatin (PF) in the treatment of unresectable esophageal cancer: a systematic review and meta-analysis of the efficacy and toxicity of two different regimens. J Gastrointest Oncol. 2023;14: 1037–1051. doi:10.21037/jgo-23-33

36. Schwaid AG, Krasowka-Zoladek A, Chi A, Cornella-Taracido I. Comparison of the Rat and Human Dorsal Root Ganglion Proteome. Sci Rep. 2018;8: 13469. doi:10.1038/s41598-018-31189-9

37. Leal-Julià M, Vilches JJ, Onieva A, Verdés S, Sánchez Á, Chillón M, et al. Proteomic quantitative study of dorsal root ganglia and sciatic nerve in type 2 diabetic mice. Mol Metab. 2021;55: 101408. doi:10.1016/j.molmet.2021.101408

38. Pogatzki-Zahn EM, Gomez-Varela D, Erdmann G, Kaschube K, Segelcke D, Schmidt M. A proteome signature for acute incisional pain in dorsal root ganglia of mice. Pain. 2021;162: 2070–2086. doi:10.1097/j.pain.0000000000002207

39. 39. Venny 2.1.0. [cited 7 Jun 2024]. Available: https://bioinfogp.cnb.csic.es/tools/venny/

40. Dengjel J, Kratchmarova I, Blagoev B. Mapping Protein–Protein Interactions by Quantitative Proteomics. In: Cutillas PR, Timms JF, editors. LC-MS/MS in Proteomics. Totowa, NJ: Humana Press; 2010. pp. 267–278. doi:10.1007/978-1-60761-780-8_16

41. de Clauser L, Kappert C, Sondermann JR, Gomez-Varela D, Flatters SJL, Schmidt M. Proteome and Network Analysis Provides Novel Insights Into Developing and Established Chemotherapy-Induced Peripheral Neuropathy. Front Pharmacol. 2022;13: 818690. doi:10.3389/fphar.2022.818690

42. Doty M, Yun S, Wang Y, Hu M, Cassidy M, Hall B, et al. Integrative multiomic analyses of dorsal root ganglia in diabetic neuropathic pain using proteomics, phospho-proteomics, and metabolomics. Sci Rep. 2022;12: 17012. doi:10.1038/s41598-022-21394-y

43. Cirrincione AM, Pellegrini AD, Dominy JR, Benjamin ME, Utkina-Sosunova I, Lotti F, et al. Paclitaxel-induced peripheral neuropathy is caused by epidermal ROS and mitochondrial damage through conserved MMP-13 activation. Sci Rep. 2020;10: 3970. doi:10.1038/s41598-020-60990-8

44. Canta A, Pozzi E, Carozzi VA. Mitochondrial Dysfunction in Chemotherapy-Induced Peripheral Neuropathy (CIPN). Toxics. 2015;3: 198–223. doi:10.3390/toxics3020198

45. Duggett NA, Griffiths LA, Flatters SJL. Paclitaxel-induced painful neuropathy is associated with changes in mitochondrial bioenergetics, glycolysis, and an energy deficit in dorsal root ganglia neurons. Pain. 2017;158: 1499–1508. doi:10.1097/j.pain.0000000000000939

46. Bennett GJ, Doyle T, Salvemini D. Mitotoxicity in distal symmetrical sensory peripheral neuropathies. Nat Rev Neurol. 2014;10: 326–336. doi:10.1038/nrneurol.2014.77

47. Huang Z-Z, Li D, Liu C-C, Cui Y, Zhu H-Q, Zhang W-W, et al. CX3CL1-mediated macrophage activation contributed to paclitaxel-induced DRG neuronal apoptosis and painful peripheral neuropathy. Brain, Behavior, and Immunity. 2014;40: 155–165. doi:10.1016/j.bbi.2014.03.014

48. Zhang H, Li Y, De Carvalho-Barbosa M, Kavelaars A, Heijnen CJ, Albrecht PJ, et al. Dorsal Root Ganglion Infiltration by Macrophages Contributes to Paclitaxel Chemotherapy-Induced Peripheral Neuropathy. The Journal of Pain. 2016;17: 775–786. doi:10.1016/j.jpain.2016.02.011

49. Woo DDL, Miao SYP, Pelayo JC, Woolf AS. Taxol inhibits progression of congenital polycystic kidney disease. Nature. 1994;368: 750–753. doi:10.1038/368750a0

50. Benbow SJ, Wozniak KM, Kulesh B, Savage A, Slusher BS, Littlefield BA, et al. Microtubule Targeting Agents Eribulin and Paclitaxel Differentially Affect Neuronal Cell Bodies in Chemotherapy Induced Peripheral Neuropathy. Neurotox Res. 2017;32: 151–162. doi:10.1007/s12640-017-9729-6

51. Flatters SJL, Bennett GJ. Studies of peripheral sensory nerves in paclitaxel-induced painful peripheral neuropathy: Evidence for mitochondrial dysfunction. Pain. 2006;122: 245–257. doi:10.1016/j.pain.2006.01.037

52. Talagas M, Lebonvallet N, Leschiera R, Marcorelles P, Misery L. What about physical contacts between epidermal keratinocytes and sensory neurons? Experimental Dermatology. 2018;27: 9–13. doi:10.1111/exd.13411

53. Nedergaard M, Ransom B, Goldman SA. New roles for astrocytes: Redefining the functional architecture of the brain. Trends in Neurosciences. 2003;26: 523–530. doi:10.1016/j.tins.2003.08.008

54. Jasmin L, VIT J-P, Bhargava A, Ohara PT. Can satellite glial cells be therapeutic targets for pain control? Neuron Glia Biol. 2010;6: 63–71. doi:10.1017/S1740925X10000098

55. Walters MC, Ladle DR. Calcium homeostasis in parvalbumin DRG neurons is altered after sciatic nerve crush and sciatic nerve transection injuries. J Neurophysiol. 2021;126: 1948– 1958. doi:10.1152/jn.00707.2020

56. Costa FAL, Moreira Neto FL. [Satellite glial cells in sensory ganglia: its role in pain]. Rev Bras Anestesiol. 2015;65: 73–81. doi:10.1016/j.bjan.2013.07.013

57. Facer P, Casula MA, Smith GD, Benham CD, Chessell IP, Bountra C, et al. Differential expression of the capsaicin receptor TRPV1 and related novel receptors TRPV3, TRPV4 and TRPM8 in normal human tissues and changes in traumatic and diabetic neuropathy. BMC Neurol. 2007;7: 11. doi:10.1186/1471-2377-7-11

58. Han Q, Kim YH, Wang X, Liu D, Zhang Z-J, Bey AL, et al. SHANK3 Deficiency Impairs Heat Hyperalgesia and TRPV1 Signaling in Primary Sensory Neurons. Neuron. 2016;92: 1279–1293. doi:10.1016/j.neuron.2016.11.007

59. Wilder-Smith CH. Abnormal endogenous pain modulation and somatic and visceral hypersensitivity in female patients with irritable bowel syndrome. WJG. 2007;13: 3699. doi:10.3748/wjg.v13.i27.3699

60. Yu X, Liu H, Hamel KA, Morvan MG, Yu S, Leff J, et al. Dorsal root ganglion macrophages contribute to both the initiation and persistence of neuropathic pain. Nat Commun. 2020;11: 264. doi:10.1038/s41467-019-13839-2

61. Klein I, Boenert J, Lange F, Christensen B, Wassermann MK, Wiesen MHJ, et al. Glia from the central and peripheral nervous system are differentially affected by paclitaxel chemotherapy via modulating their neuroinflammatory and neuroregenerative properties. Front Pharmacol. 2022;13: 1038285. doi:10.3389/fphar.2022.1038285

62. Huang T-Y, Belzer V, Hanani M. Gap junctions in dorsal root ganglia: Possible contribution to visceral pain. European Journal of Pain. 2010;14: 49.e1–49.e9. doi:10.1016/j.ejpain.2009.02.005

63. Warwick R a., Hanani M. The contribution of satellite glial cells to chemotherapy-induced neuropathic pain. European Journal of Pain. 2013;17: 571–580. doi:10.1002/j.1532-2149.2012.00219.x

64. Suadicani SO, Cherkas PS, Zuckerman J, Smith DN, Spray DC, Hanani M. Bidirectional calcium signaling between satellite glial cells and neurons in cultured mouse trigeminal ganglia. Neuron Glia Biol. 2010;6: 43–51. doi:10.1017/S1740925X09990408

65. Kushnir R, Cherkas PS, Hanani M. Peripheral inflammation upregulates P2X receptor expression in satellite glial cells of mouse trigeminal ganglia: A calcium imaging study. Neuropharmacology. 2011;61: 739–746. doi:10.1016/j.neuropharm.2011.05.019

66. Mikesell AR, Isaeva E, Schulte ML, Menzel AD, Sriram A, Prahl MM, et al. Keratinocyte Piezo1 drives paclitaxel-induced mechanical hypersensitivity. bioRxiv. 2023; 2023.12.12.571332. doi:10.1101/2023.12.12.571332

67. Komori N, Takemori N, Kim HK, Singh A, Hwang S-H, Foreman RD, et al. Proteomics study of neuropathic and nonneuropathic dorsal root ganglia: altered protein regulation following segmental spinal nerve ligation injury. Physiological Genomics. 2007;29: 215–230. doi:10.1152/physiolgenomics.00255.2006

68. Gerdle B, Ghafouri B. Proteomic studies of common chronic pain conditions - a systematic review and associated network analyses. Expert Review of Proteomics. 2020;17: 483–505. doi:10.1080/14789450.2020.1797499

69. Ahlawat A, Rana A, Goyal N, Sharma S. Potential role of nitric oxide synthase isoforms in pathophysiology of neuropathic pain. Inflammopharmacology. 2014;22: 269–278. doi:10.1007/s10787-014-0213-0

70. Hara MR, Snyder SH. Cell Signaling and Neuronal Death. Annual Review of Pharmacology and Toxicology. 2007;47: 117–141. doi:10.1146/annurev.pharmtox.47.120505.105311

71. Hamza M, Wang X-M, Wu T, Brahim JS, Rowan JS, Dionne RA. Nitric oxide is negatively correlated to pain during acute inflammation. Mol Pain. 2010;6: 55. doi:10.1186/1744-8069-6-55

72. Peng X, Studholme K, Kanjiya MP, Luk J, Bogdan D, Elmes MW, et al. Fatty-acid-binding protein inhibition produces analgesic effects through peripheral and central mechanisms. Mol Pain. 2017;13: 1744806917697007. doi:10.1177/1744806917697007

